# G4mer: An RNA language model for transcriptome-wide identification of G-quadruplexes and disease variants from population-scale genetic data

**DOI:** 10.1101/2024.10.01.616124

**Authors:** Farica Zhuang, Danielle Gutman, Nathaniel Islas, Bryan B Guzman, Alli Jimenez, San Jewell, Nicholas J Hand, Katherine Nathanson, Daniel Dominguez, Yoseph Barash

## Abstract

RNA G-quadruplexes (rG4s) are key regulatory elements in gene expression, yet the effects of genetic variants on rG4 formation remain underexplored. Here, we introduce G4mer, an RNA language model that predicts rG4 formation and evaluates the effects of genetic variants across the transcriptome. G4mer significantly improves accuracy over existing methods, highlighting sequence length and flanking motifs as important rG4 features. Applying G4mer to 5’ untranslated region (UTR) variations, we identify variants in breast cancer-associated genes that alter rG4 formation and validate their impact on structure and gene expression. These results demonstrate the potential of integrating computational models with experimental approaches to study rG4 function, especially in diseases where non-coding variants are often overlooked. To support broader applications, G4mer is available as both a web tool and a downloadable model.

## Introduction

RNA G-quadruplexes (rG4s) are secondary structures formed in guanine-rich regions of RNA, which have emerged as crucial regulatory elements in gene expression. These structure motifs are particularly enriched in UTRs of mRNAs^1,2^, where they play key roles in various regulatory processes including translation, alternative splicing, polyadenylation, and mRNA stability^3–5^. Alterations to rG4 structures have been linked to human pathologies, such as cancer and neurological diseases, by disrupting the gene expression landscape in affected individuals^6^. Furthermore, the effects of rG4 alterations on functions like translation have been investigated, revealing that changes in rG4 formation can significantly impact these processes^7–9^. Consequently, rG4s have garnered increasing attention for their therapeutic potential^10–13^. Despite the growing recognition of their importance, the influence of genetic variants on rG4 formation, particularly at the transcriptome-wide level, remains poorly understood.

Recent advancements in high-throughput sequencing, such as rG4-seq^1^, have enabled the mapping of rG4s across the transcriptome. However, these methods are often limited by noise, condition specificity, and the inability to capture rG4s in lowly expressed genes^14^. As a result, computational models to predict G4 formation have been developed. Traditional methods are score-based, typically relying on G/C content, skewness, and the number of consecutive Gs present in the sequence to score the propensity of G4s^15–19^. However, such methods may overlook noncanonical forms and struggle in regions where flanking sequences introduce noise by altering the G/C ratio, leading to inaccurate predictions of rG4 formation. More recently, machine learning-based models have been developed to predict rG4 formation^20–22^. These models, particularly those based on convolutional neural networks (CNNs)^23–26^, have achieved state-of-the-art performance in predicting rG4 structures by learning patterns from labeled datasets. However, CNN-based models are inherently limited by the size of their convolutional kernels, which are designed to capture short-range interactions. This limitation may make it challenging for CNNs to accurately predict rG4 formation in longer, transcriptome-wide sequences. Arguably, a more accurate model can not only improve predictions of rG4 formation, but also help detect the effects of genetic variants affecting rG4s as well as gain insights into the sequence features governing rG4 formation.

To create such a model, we developed G4mer, a transformer-based^27^ RNA language model designed to predict rG4 formations across the human transcriptome. We first evaluated its performance against current state-of-the-art methods across diverse data and experimental techniques, showing that it significantly outperforms existing methods. We then leveraged G4mer to analyze the impact of transcriptome-wide genetic variants on rG4s, focusing particularly on the 5’ and 3’ UTRs of mRNAs where rG4s are most densely populated^1,2^, and assessed negative selection of rG4-disrupting variants. Our analyses revealed that rG4 length is an overlooked but potentially crucial factor in rG4 prediction, experimental analysis, and negative selection. Additionally, we explored disease associations of rG4-altering variants from the Penn Medicine Biobank (PMBB)^28^ that are predicted to disrupt or stabilize rG4 structures. We demonstrate how G4mer can be applied with phenome-wide association study (PheWAS) to identify rG4-altering variants in breast cancer-associated genes. We then show that the variants we identified also impact structure formation using circular dichroism spectroscopy and validated the functional effects of these rG4-altering variants on protein expression using dual luciferase reporter assays, linking rG4 structural changes with functional outcomes. Finally, to facilitate research of other rG4 and genetic variants that may affect those, we create a web tool that allows users to predict and download rG4 predictions for transcriptomic sequences.

## Results

### G4mer is a transformer-based model that improves rG4 formation and subtype predictions

To develop G4mer, we first pre-trained mRNAbert, a bidirectional encoder representation model (BERT)^29^, using masked language modeling on the entire human transcriptome (see Methods). The pre-training step allowed the model to learn generalized representations of RNA sequences in a self-supervised manner. Following pre-training, we fine-tuned mRNAbert specifically for rG4 prediction using a high-throughput rG4 detection experimental dataset^30^ along with non-rG4 sequences (Fig.1b). For this, we utilized the rG4-seeker dataset, which contains over 5000 regions identified as containing rG4 elements through the rG4-seq^1^ in HeLa cells, then post-processed to ensure accurate detection of rG4 sequences^30^. To complement this, an equal number of G-rich negative sequences, sampled from transcripts with no rG4 detected, were included to define the non-rG4 sequences (see Methods for additional details). We first performed standard 10-fold cross-validation (CV) on the dataset of rG4 and non-rG4 sequences to evaluate rG4 prediction accuracy of G4mer compared to the CNN-based model G4detector^23^, which previously achieved state-of-the-art performance in predicting G4 formation. On this task, G4mer outperformed G4Detector, obtaining an accuracy of 0.95 and a receiver operating characteristic-area under the curve (ROC-AUC) score of 0.98, compared to an accuracy of 0.92 and an AUC of 0.97 for G4Detector (Fig.1c).

**Figure 1.**
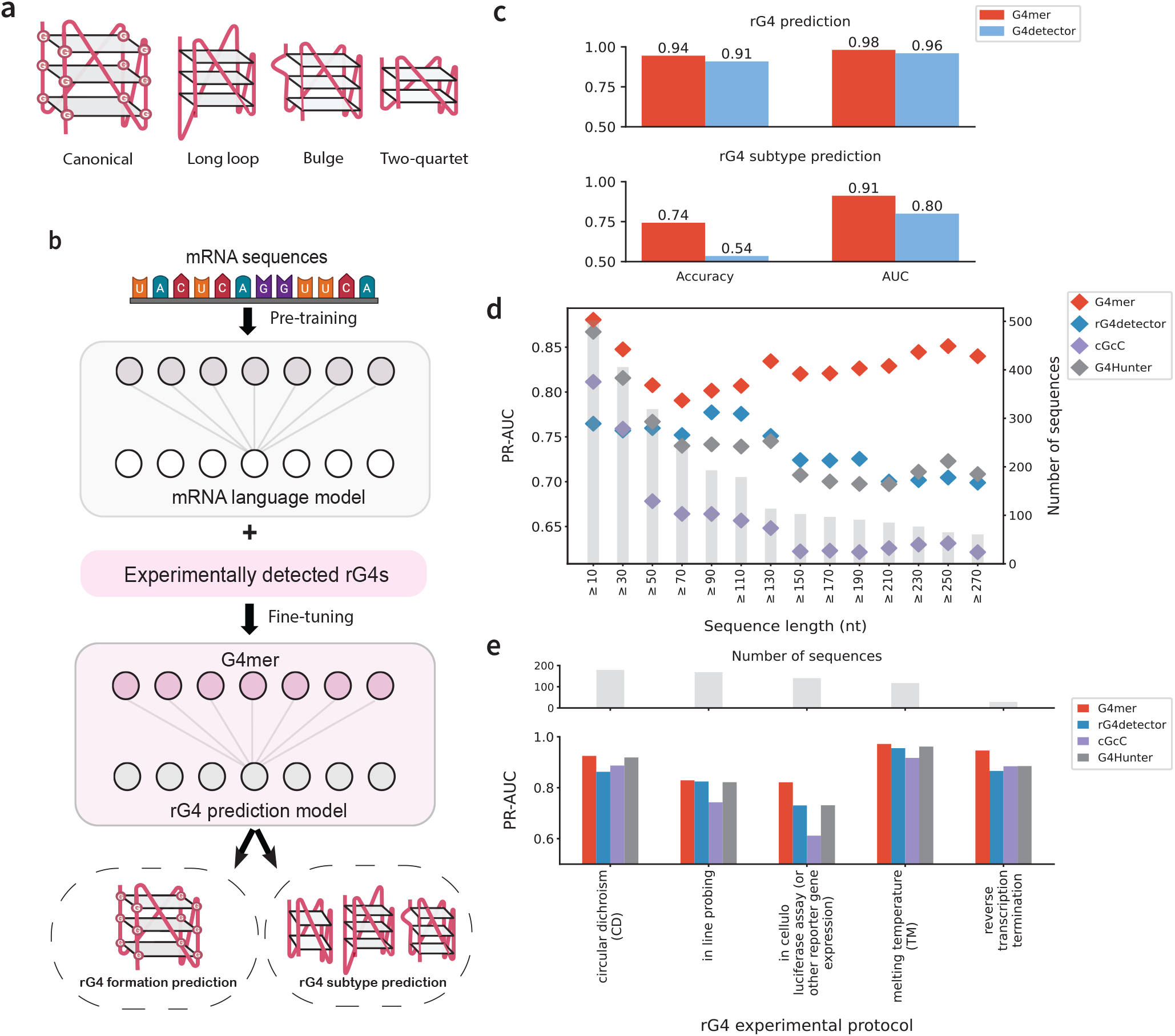
**(a)** rG4 structures can be categorized into canonical as well as noncanonical subtypes such as long loop, bulge, and two-quartet. **(b)** G4mer is developed based on an mRNA language model that was pre-trained on the entire human transcriptome and fine-tuned with experimentally detected rG4 sequences. **(c)** Comparison of transformer-based G4mer (red) and CNN-based rG4detector (blue) for rG4 binary prediction (top) and rG4 subtype multiclass prediction (bottom). Performance is measured by accuracy and ROC-AUC. **(d)** PR-AUC performance of G4mer compared to other models (rG4detector, cGcC, G4Hunter) on sequences of increasing lengths from the G4RNA database. The gray bars represent the number of sequences available in each length category, while the colored points indicate the PR-AUC for each model as sequence lengths increase. **(e)** PR-AUC comparison of G4mer and other models across the top 5 experimental protocols for rG4 detection, ranked by the highest number of sequences reported in the G4RNA database. The bar plot at the top shows the number of sequences for each protocol, while the lower plot compares the predictive performance (PR-AUC) of G4mer, rG4detector, cGcC, and G4Hunter across these protocols.

Following this initial comparison, we investigated the models’ abilities in predicting rG4 subtypes on the aforementioned rG4-seeker dataset. In this data, the authors categorized each sequence into one of eight labels, ranging from canonical to known noncanonical subtypes such as long loops, bulges, and two-quartets. Other categories include different G-rich sequence patterns and a category for sequences that do not fall into any of the defined groups. rG4 sequences that matched the pattern of multiple categories were assigned to the class with the higher predicted stability, following the order of canonical rG4s, long loops, bulges, two-quartets, and so on^1^. Therefore, identifying the rG4 subtype category of a given sequence requires a model to learn not only the sequence patterns and logical rules but also the associated stability of the sequences. In this context, G4mer significantly outperformed G4Detector, achieving an accuracy of 0.80 and an AUC of 0.92, compared to an accuracy of 0.55 and an AUC of 0.79 for G4Detector (Fig.1c). G4mer’s improved subtype classification demonstrates its ability to capture intricate sequence patterns and implicitly assess the stability of the sequences. Overall, these results highlight the robustness of G4mer’s transformer-based architecture and its superior predictive capability in identifying rG4 formation, both in general and for specific subtypes.

We next sought to assess the prediction capabilities beyond the specific data and experimental techniques of rG4-seq. We therefore turned to an independent set of 795 published sequences from the G4RNA database^31^. The G4RNA database includes sequences tested under various experimental protocols from different publications (Supplementary Fig. S2a). The sequences were filtered to ensure consistency, including the removal of sequences reported by multiple sources with conflicting labels (Methods). We then compared G4mer’s performance against CNN-based rG4detector^24^, as well as the scoring-based methods cGcC^15^ and G4Hunter^16^. G4mer outperformed all the models across all the filtered G4RNA sequences (Supplementary Fig. S1).

Notably, the G4RNA database includes artificial sequences, transcript segments, and full-length UTR sequences, resulting in lengths ranging from 14 to 1,368 nt. Although flanking regions are important for accurate rG4 formation prediction, including excessive lengths in the sequence input may introduce noise, for example, by adding additional G-rich elements that can confuse the model and reduce prediction accuracy. When evaluating all sequences (10 nt or longer), G4Hunter performance was close to that of G4mer (Fig.1d). However, as we focused on progressively longer sequences, a significant performance difference emerged. G4mer remained less sensitive to increasing sequence length, consistently outperforming other models with PR-AUC values around 0.85 for all lengths. In contrast, other models suffered from a noticeable decline in performance (Fig.1d). This suggests G4mer’s robustness in handling longer sequences, making it suitable for transcript-wide prediction analyses, particularly those involving UTR regions that span hundreds to thousands of nucleotides.

To assess G4mer’s generalizability to sequences from diverse experimental protocols, we compared the performance of G4mer, G4detector, cGcC, and G4Hunter across the top 5 rG4 experimental protocols in G4RNA (Fig 1e). These protocols included circular dichroism (CD), in-line probing, in cellulo luciferase assay (or other reporter gene expression), melting temperature (Tm), and reverse transcription termination. G4mer offers improved performance for sequences across all experimental protocols. Notably, G4mer performs well on sequences from both the CD protocol and the in cellulo luciferase assay, achieving a PR-AUC of 0.82 despite the stark differences in their sequence length distributions (Supplementary Fig. S2b). Specifically, the CD protocol has a median sequence length of 30.5 nt, while the in cellulo luciferase assay has a median of 210 nt (Supplementary Fig. S2c). This demonstrates G4mer’s robust ability to accurately predict rG4 formation across diverse experimental conditions, even when faced with varying sequence length distributions that arise from technical differences. These results underscore that, despite being trained on rG4-seq protocol data, G4mer successfully captures the fundamental properties of rG4s, allowing it to generalize well and minimize the impact of technical biases.

While the performance comparison highlights the accuracy of rG4 predictions across the transcriptome, it raises the question of what features, beyond the G-tracts themselves, may be contributing to G4mer’s improved predictions. To address this question, we analyzed the flanking regions of predicted rG4 sequences in UTRs across the transcriptome using Enhanced Integrated Gradient (EIG) analysis^32^, a method for deep learning model interpretation. EIG quantifies the contribution of each feature, specifically k-mers in this context, to a model’s prediction by integrating the gradient along a path from a reference point to a sample of interest. Using EIG, we obtained attribution scores for k-mers of different lengths in the flanking regions of rG4 sequences, identifying those that significantly contributed to the prediction compared to non-rG4 sequences (*p <* 0.05, two-sided t-test, Benjamini-Hochberg FDR-adjusted, see Methods for details). The EIG analysis shows that the sequences flanking the G-tracts do indeed contribute to the rG4 predictions (Supplementary Fig SS3a). Specifically, we find that the positions proximal to the G-tracts contribute more to the prediction and that guanine and uracil are preferred in those positions, while cytosine bases generally contributed negatively to predicting rG4 formation. However, the EIG analysis clearly indicates that analyzing attribution per position independently, as is often done when constructing position-specific scoring motifs (PSSM), can be misleading. For example, some cytosines near rG4 motifs showed positive attribution scores, but we observed distinct attribution score patterns for 2-mers and 3-mers (Supplementary Fig. S3b,c). CC dinucleotides consistently had negative attribution scores, whereas dinucleotides containing a cytosine and another nucleotide could show positive attributions. Similarly, trinucleotides with predominantly cytosine bases were negatively attributed unless other nucleotides were present. In contrast, UU dinucleotides had a strong positive influence on rG4 predictions. Overall, the above results suggest that rG4 formation, as captured by G4mer, avoids competing G-C bounds and prefers open regions via stretches of uridine, emphasizing the role of flanking regions in rG4 formation.

### G4mer allows transcriptome-wide mapping of rG4-altering variants to assess their functional significance

Having established the high accuracy of G4mer in predicting rG4 formation given a sequence, we leveraged its capabilities to predict the effects of single nucleotide variants (SNVs) on rG4 formation probability transcriptome-wide. Specifically, rG4-altering variants that significantly lower the rG4 prediction probability (from > 0.7 to < 0.3, see Methods) are considered rG4-breaking. We analyzed the entire set of genetic variants from the gnomAD database^33^, focusing on subgroups of rG4-breaking and non-breaking variants to detect signals of negative selection.

Negative selection, where deleterious variants are continuously removed from the population, is crucial for maintaining biological function. Signals of negative selection can be identified by the depletion of variants or, conversely, by an unexpected enrichment of rare variants within a specific class. Therefore, to detect negative selection in rG4 elements, we first evaluated the mean alternate allele frequencies (MAFs) of variants predicted to break rG4s (rG4-breaking) and those in rG4s that do not disrupt the structure (rG4 non-breaking) from regions of matched genomic constraints. Genomic constraint was quantified using the constraint Z-score, which reflects the deviation of observed from expected variations in a given genomic region^33^. To account for differences in sequencing depth in the gnomAD whole genome sequencing dataset, we excluded variants with a call rate below 80%. Overall, we found 8,785 5’UTR and 17,181 3’UTR variants in gnomAD that are rG4-breaking. Conversely, we found 18,049 5’UTR variants and 30,005 3’UTR variants in rG4s that have negligible effects on rG4 structure prediction in similarly constrained regions. Notably, rG4-breaking variants show a significant reduction in mean allele frequency compared to variants in rG4s that don’t affect the structure (p « 4.24e-16, one-tailed KS test) (Fig. 2a). Interestingly, the subset of these variants that break the consecutive G-run section of the rG4 sequences with similar nucleotide contexts also shows a significant reduction in mean allele frequencies for rG4-breaking variants, consistent with the effects of negative selection. Finally, additional analysis revealed a positive correlation between the magnitude of rG4 disruption (more negative G4mer delta scores) and higher Combined Annotation Dependent Depletion (CADD) scores, particularly for variants in the 5’UTR region (Fig.2b). This relationship suggests that variants that significantly disrupt rG4 structures are more likely to be deleterious, supporting the hypothesis that rG4s may play a critical functional role.

**Figure 2.**
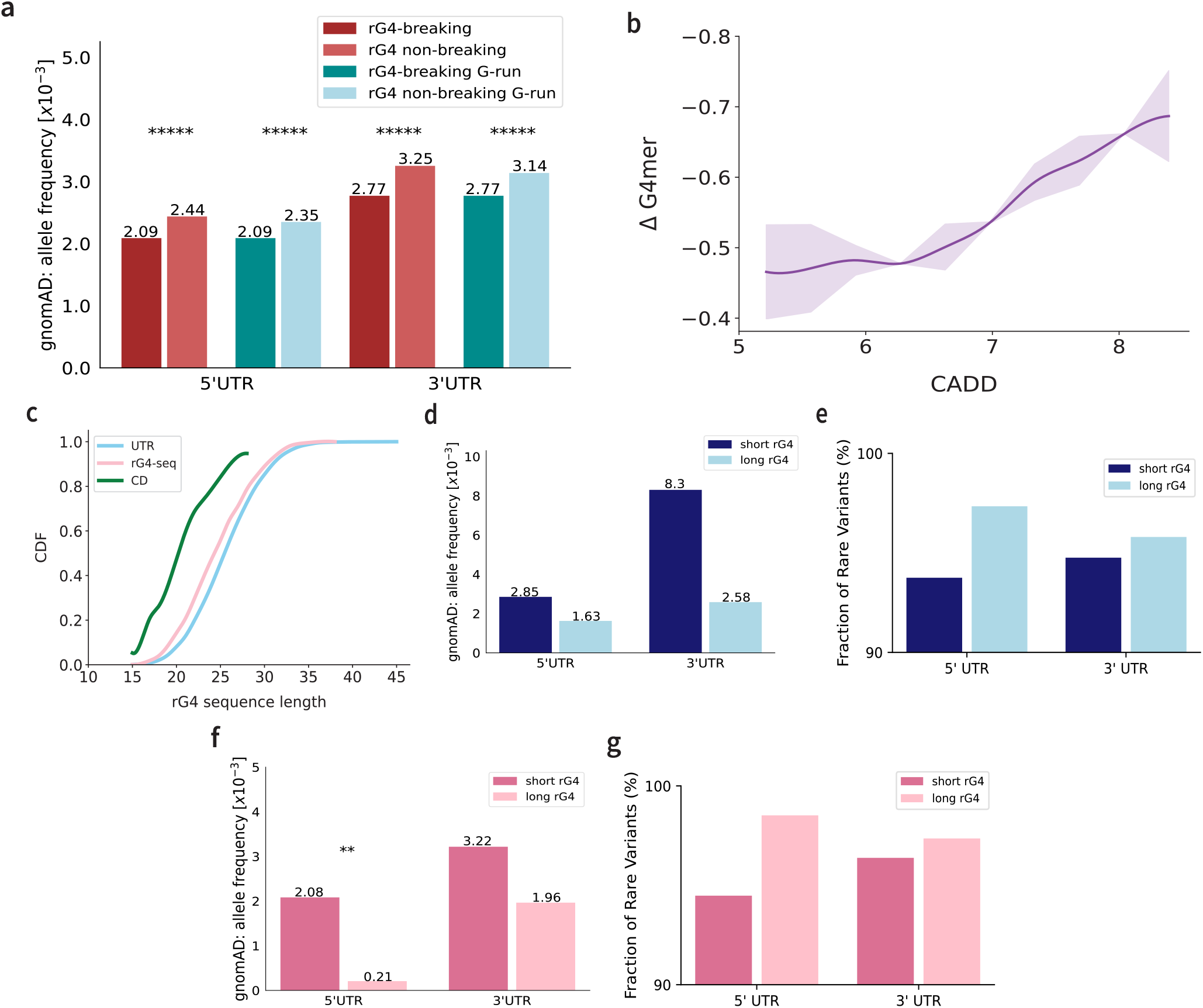
rG4-altering variant analyses. **(a)** gnomAD mean alternate allele frequency comparison between rG4-breaking variants and negative control rG4 variants that are non-breaking across the 5’UTR and 3’UTR in all contexts (red) as well as in the G-runs (blue). Asterisks denote significance (p « 4.24e-16, one-tailed KS test). **(b)** Relationship between raw CADD scores and ΔG4mer values in 5’UTR regions. ΔG4mer represents the change in predicted rG4 formation probability when comparing sequences with and without a variant. **(c)** Cumulative distribution function (CDF) plot comparing the length distributions of putative canonical rG4s identified in UTR regions (blue), sequences from the rG4-seq experiment conducted on HeLa cells (pink), and sequences that were validated using circular dichroism (CD) from the G4RNA database (green). **(d)** Mean alternate allele frequency comparisons of variants breaking short versus long canonical rG4. **(e)** Fraction of rare variants (MAF < 0.1%) breaking short versus long canonical rG4. **(f, g)** Comparison of mean alternate allele frequency and fraction of rare variants for variants breaking canonical and long-loop rG4s. Asterisks denote significance (p < 0.028, one-tailed Mann-Whitney U test).

Building on the variant analyses above, which underscore the functional significance of rG4s, we next aimed to explore whether the importance of rG4s varies depending on their sequence characteristics, such as length. Specifically, our previous results (Fig. 1d, Supplementary Fig. S2) revealed a strong bias in some existing models and experimental protocols for short rG4 sequences. This performance variation raises the question of whether such a bias may, in turn, lead to overlooking longer rG4s that have functional significance. Naturally, rG4 subtypes could greatly affect the sequence length. For example, the noncanonical long loops are intrinsically longer than canonical rG4s. To minimize variations in lengths contributed by rG4 subtypes, we first focused on canonical rG4s. We observed that the length distributions of sequences could vary based on the experimental technique used. For instance, rG4-seq tends to capture longer rG4 sequences with a median of 24 nt, compared to a commonly used structure validation method, circular dichroism spectroscopy, with a median of 20 nt (Fig. 2c). Notably, putative rG4s in the UTR regions of the human transcriptome often consist of longer sequences with a median of 25 nt that more closely resemble those detected by the rG4-seq protocol. Moreover, tested sequences with CD have a maximum length of 28 nt, while the longest putative rG4s found in the UTR span 45 nt.

Next, we investigated the MAF of variants located within short (≤22 nt) and long (28 nt≤) rG4 sequences across both 5’UTR and 3’UTR regions. Distinct differences emerged between the variants located in short and long rG4 sequences across these regions (Figure 2d). Specifically, the mean allele frequency for constraint-matched variants breaking long canonical rG4 sequences is lower compared to those within short rG4 sequences. Additionally, we observed a similar trend where long canonical rG4 sequences showed a higher fraction of rare variants (MAF < 0.1%) (Fig. 2e). These trends, though not statistically significant with the limited number of variants tested, suggest that long rG4s may be equally, if not more, functionally important than short rG4s in the transcriptome. Finally, to expand beyond canonical rG4s, we included the noncanonical long loop rG4s and increased the definition of long rG4s (≥ 31 nt). This larger gap between short and long rG4 definitions allowed us to observe a significant reduction in the mean allele frequency of variants breaking long rG4s (p < 0.028, one-tailed Mann-Whitney U test) (Fig. 2g). Moreover, we noticed a similar trend in which variants breaking long rG4s exhibited a higher fraction of rare variants across the UTR regions (Fig. 2h).

### Non-coding rG4-altering variants are associated with breast cancer and modulate gene expression

The reduced MAF observed in rG4-altering variants suggests these regions may have functional significance and could play a role in human diseases. To explore the potential contribution of rG4-altering variants to disease etiology, we focused on breast cancer, a condition where much of the disease heritability remains unexplained^34–36^. Specifically, we performed two types of analysis. First, we performed PheWAS for breast cancer using data from the Penn Medicine BioBank (PMBB, see Methods). Second, we evaluated PMBB variants in 11 genes already implicated in breast cancer for their potential to create or disrupt rG4s in the respective UTR regions. Overall, we identified 28 rG4-breaking variants and 29 rG4-forming variants across 8 genes. The full list of variants is included in Supplementary Table 1, with key findings highlighted below.

We detected a significant PheWAS association (p = 7.4e-7) between an rG4-breaking variant in the 5’ UTR of *EPN3* and breast cancer (Fig.3a). Although *EPN3* was not included in our initial set of breast cancer-associated genes, previous studies have identified it as an oncogene involved in breast cancer by regulating epithelial-mesenchymal transition, playing a crucial role in metastasis^37^. Furthermore, *EPN3* has been explored as a potential therapeutic target due to its role in regulating apoptosis in breast cancer cells^38^. Hence, the association we observed suggests that rG4-breaking variants may have a broader impact on disease phenotypes and indicate that rG4 structures within the 5’UTR of *EPN3* may play a critical role in gene regulation. The disruption of rG4s could potentially alter downstream effects, such as gene expression, protein function, or interaction with other cellular components, thereby contributing to cancer susceptibility. Another notable rG4-altering variant was detected in DNA mismatch repair gene *MSH6*. In this case, the variant was predicted to form, rather than disrupt, an rG4 in the 5’UTR. *MSH6* has been extensively studied for its associations with Lynch syndrome and its role in increasing the risk of several cancers^39–42^.

**Figure 3.**
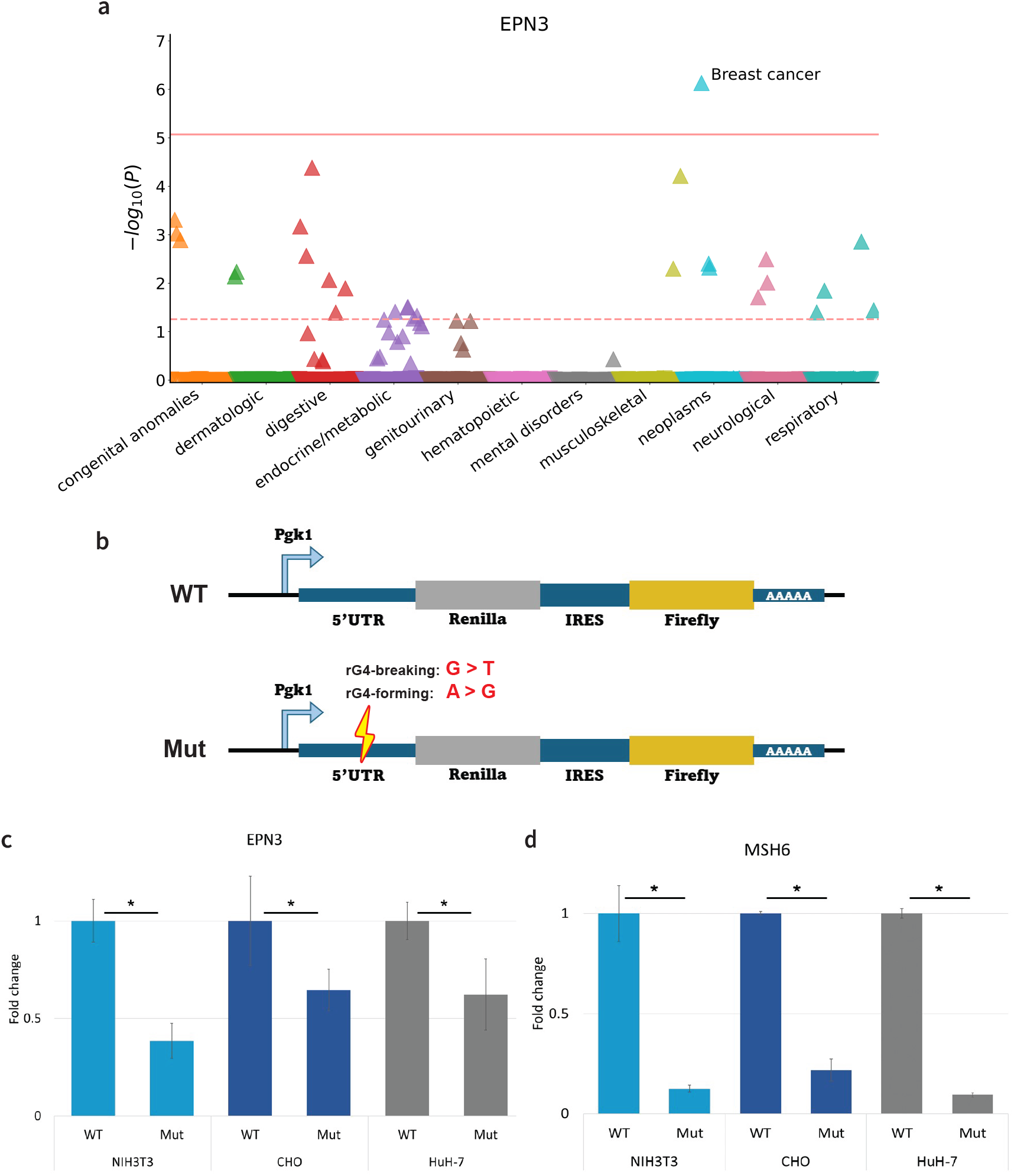
Disease-associated rG4-altering variants change protein expression level. **(a)** PheWAS plot of rG4-disrupting variant in the 5’UTR of *EPN3* (N=1,201 carriers) in the Penn Medicine BioBank. Each point represents a single Phecode based mapped from ICD-10 and plotted by disease category on the x-axis. The height of each point corresponds to the −log_10_(P value) of the association between the variant and phenotype tested using a logistic regression model adjusted for age at enrollment, age^2^, sex, and the first ten principal components (PCs) of genetic ancestry. The solid red line represents the threshold for Bonferroni-adjusted significance (P=8.5e-6) and the red dashed line represents the FDR < 0.1 threshold (P=5.5e-2) accounting for multiple comparisons. The direction of each arrowhead corresponds to increased risk (up) or decreased risk (down) represented by odds ratio. **(b)** Schematic representation of the wildtype (WT) and mutant (Mut) dual luciferase reporter constructs used to study the functional effects of rG4 formation in the 5’UTR. The WT construct contains the native 5’UTR, while the Mut construct includes single nucleotide mutations in the 5’UTR. The mutations tested are rG4-breaking (G>T) or rG4-forming (A>G), which are shown in red. Renilla and Firefly luciferase activities are measured to assess the functional impact of these mutations on rG4 stability and translation efficiency. **(c, d)** Dual-luciferase reporter assay quantifies relative expression for (c) rG4-disrupting variant in *EPN3* and (d) rG4-forming variant in *MSH6*. Mean ±SEM of relative luciferase activity shown, statistical significance is denoted by asterisks (* p < 1e-4, independent t-test) for 3-5 independent experiments with technical repeats across three different cell lines (NIH3T3, CHO, and HuH-7).

Given the established role of rG4s within the 5’UTR in translational regulation^3,7,43^, we sought to investigate the functional impact of the aforementioned rG4-altering variants using dual luciferase reporter assays across three distinct cell lines: NIH3T3, HuH-7, and CHO. For each gene, we constructed two 5’UTR constructs: the wild-type (WT) sequence and a mutated (Mut) sequence designed to alter the rG4 formation (Fig. 3b, See Methods). Our results demonstrated a significant modulation of protein expression by rG4-altering variants in both *EPN3* and *MSH6* across all tested cell lines. Specifically, the presence of disease-associated rG4-breaking variants led to a marked reduction in protein expression compared to the wild-type sequences in the case of *EPN3* in all three cell lines (Fig.3c). Similarly, the rG4-forming variant in *MSH6* consistently resulted in a significant downregulation of protein expression across all cell lines, highlighting the regulatory roles that rG4 structures may play in gene expression (Fig.3d). Overall, the rG4-altering single nucleotide variants predicted by G4mer induced significant changes to protein expression levels.

### Structural validation of predicted rG4-altering variants with circular dichroism spectroscopy

The observed changes in protein expression due to the predicted rG4-altering variants indicate their functional effects. To explore whether these functional changes were linked to the predicted alterations in rG4 formation, we conducted structural validation using circular dichroism (CD) spectroscopy on RNA sequences from *EPN3* and *MSH6*, both with and without the rG4-altering variants predicted by G4mer. Specifically, we tested the predicted rG4-breaking variant in the *EPN3* and the predicted rG4-forming variant in the *MSH6* RNA sequence (Supplementary Table 2).

Circular dichroism spectroscopy is used to detect the presence and stability of rG4 structures by measuring the differential absorption of left- and right-handed circularly polarized light^44^. G4 structures that fold intramolecularly in a parallel topology, where the G-tracts run in the same direction, typically exhibit characteristic CD spectra with a slightly negative peak measured in millidegrees (mdeg) at 240nm and a positive peak around 260 nm^45^ when stabilized by potassium ions (KCl), as potassium specifically supports G4 formation^1,46–48^. Conversely, lithium ions (LiCl) are known to destabilize G4 structures, making them less likely to form or maintain stability under these conditions^1,49,50^. Temperature also plays a critical role; as it is increased from 25°C to 95°C, the thermal energy can create denaturing conditions that disrupt rG4 structures, resulting in a decrease in the characteristic ellipticity at 260 nm^1,51–54^.

First, we measured the CD spectra of the wild-type (WT) and mutant *EPN3* RNA under various conditions: 150 mM KCl or 150 mM LiCl at both 25°C and 95°C. In the rG4-stabilizing KCl buffer at 25°C, the WT *EPN3* RNA with a predicted rG4 displayed a characteristic positive peak at 260 nm, indicating a stable rG4 structure (Fig.4a). In contrast, the mutated RNA exhibited a significantly reduced ellipticity at 260 nm (p < 0.01), suggesting that the predicted rG4-breaking variant effectively disrupted the rG4 structure. Under the rG4-destabilizing LiCl buffer at 25°C, the ellipticity of the WT sequence decreased, reaching levels similar to the predicted disrupted rG4 in the mutated sequence. Notably, the mutated RNA maintained a similar ellipticity in both KCl and LiCl buffers, reinforcing the conclusion that the mutant is indeed a disrupted rG4 structure, unaffected by the different rG4 stabilizing effects of the buffers. At 95°C, the spectra of both WT and mutant RNAs exhibited a further decrease in ellipticity, consistent with the thermal destabilization of the rG4 structure. Thus, these results validated the rG4 breaking effect of the variant in *EPN3*.

**Figure 4.**
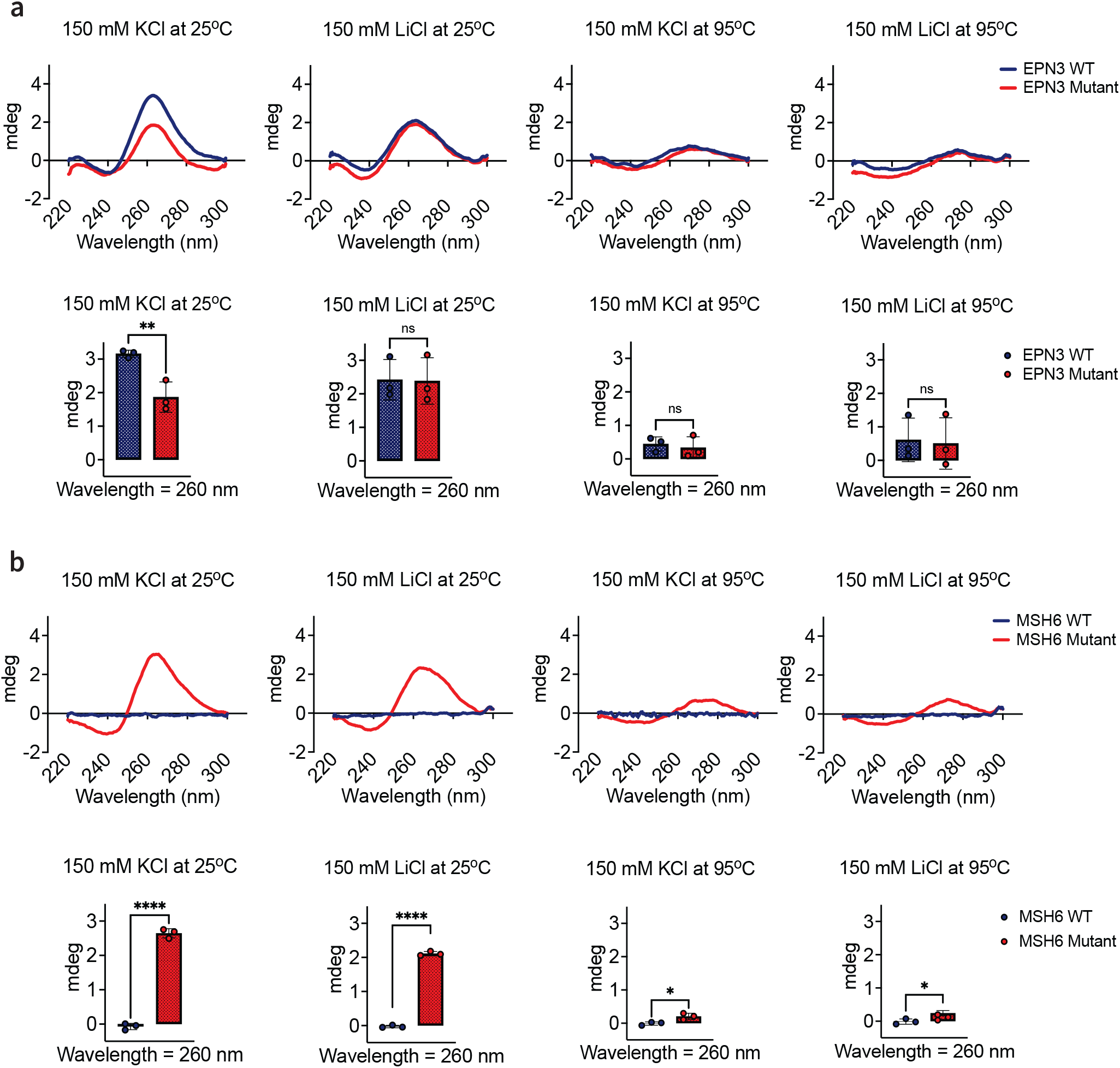
Circular dichroism (CD) spectra validates the predicted rG4-altering effects of variants in *EPN3* and *MSH6*. **(a)** CD spectra measured in ellipticity in millidegrees (mdeg) of the predicted rG4-breaking variant in the *EPN3* RNA sequence (red), compared to the wild-type (WT) (blue), under different buffer conditions (150 mM KCl or 150 mM LiCl) and temperatures (25°C and 95°C). **(b)** CD spectra of the predicted rG4-forming variant in the *MSH6* RNA sequence (red), compared to the WT (blue), under the same conditions. Each bar plot shows the mean CD signals for the WT and mutated sequences at 260nm, with error bars representing the standard deviation of three independent replicates per condition. Statistical significance is denoted by asterisks (* p < 0.05, ** p < 0.01, **** p < 1e-4, independent t-test).

Next, we measured the CD spectra for the *MSH6* RNA sequence under the same conditions. The spectra of the predicted rG4-forming variant showed an ellipticity at 260 nm that is characteristic of an rG4 under the stabilizing KCl conditions at 25°C (Fig.4b). Notably, the WT *MSH6* RNA, which was predicted to not form an rG4, showed a significantly lower ellipticity value with little to no signal at 260 nm (p < 1e-4, independent t-test), suggesting the absence of a stable rG4 structure. In the destabilizing LiCl buffer at the same temperature of 25°C, the mutated sequence with the predicted rG4-forming variant showed a decrease in ellipticity, suggesting that the predicted rG4 formation was destabilized. There was further reduction in signals at 95°C under both buffer conditions, where rG4 structures are expected to be less stable. In all conditions, the WT sequence with no predicted rG4 consistently showed little to no changes in ellipticity, confirming the absence of rG4 in the sequence. Hence, this finding supports G4mer’s prediction that the *MSH6* variant promotes rG4 formation in the mutated sequence.

Overall, our structure probing results using CD validate the predictions of G4mer, demonstrating that the *EPN3* variant disrupts an existing rG4 structure, whereby the *MSH6* variant induces rG4 formation, suggesting a link between those structural changes and the observed changes in the downstream gene protein levels measured by the dual luciferase assay.

## Discussion

We introduced G4mer, an RNA language model for predicting rG4s, built on the mRNAbert model trained on the entire human transcriptome. The pre-trained mRNAbert foundation model allows for flexible scaling of the training dataset during the fine-tuning phase to develop G4mer. Leveraging high-throughput rG4-seq data, G4mer outperformed existing state-of-the-art models and showed greater resilience to variations in input sequence lengths. As a transformer-based model, G4mer provides significant performance gains with longer input sequences, which can be attributed to the transformer model’s ability to capture long-range dependencies in RNA structures. Through model interpretation with EIG, we found guanines in the flanking regions of the rG4 motif to positively attribute to rG4 prediction. This is consistent with previous studies on DNA G4s which identified the importance of flanking regions, particularly additional G-tracts, in G4 stability^20^. Hence, G4mer is ideal for transcriptome-wide rG4 predictions on full-length transcripts spanning hundreds to thousands of nucleotides and translates well to in situ experiments. It also maintains strong results on shorter sequences and in vitro experiments. This work highlights the generalizability of G4mer across a range of rG4 sequence lengths and experimental protocols, suggesting that G4mer can be applied to both in situ and in vitro studies of RNA G-quadruplex formations, with potential applications in RNA-targeted therapies.

While existing G4 prediction models have been utilized to study the effects of variants on rG4 stability^24,55^, we are the first, to the best of our knowledge, to study transcriptome-wide effects of genetic variants on rG4 structures. Using G4mer to study the transcriptome-wide effects of variants on rG4 structure formation, G4mer was able to capture the sequence changes that affect rG4 structure stability. Moreover, variants that were predicted by G4mer to significantly break rG4s showed to be under negative selection, consistent with a previous work studying putative canonical rG4s^56^. One notable finding from our analyses of prediction models, experimental datasets, and human UTRs is the effect of rG4 length. While short rG4 sequences have been recognized for their stability^8^ and have been the focus of many experimental works, our findings suggest that longer rG4 sequences are less represented in both experimental data and prediction models, but may be at least equally functionally important with heightened negative selection and warrant further investigation. One possible reason for the heightened negative selection in longer rG4 may be that the formed structure is weaker and more flexible, allowing a single nucleotide variation to exert strong effects on their structure or the binding by RNA binding proteins (RBPs) that target these regions. Such increased volatility to genetic variants has been observed in other elements of RNA processing such as splicing^57^, and warrant further investigation in the context of rG4s.

G4mer’s application extends beyond merely predicting rG4 formations; it also provides insights into the functional and structural consequences of genetic variants. We found that rG4-altering variants could be associated with diseases. Specifically, in the disease-associated *EPN3* and *MSH6* genes, the respective rG4-breaking and rG4-forming variants led to marked reductions in protein expression compared to the wild-type sequences across multiple cell lines, suggesting that rG4 disruption by genetic variants may contribute to disease phenotypes through altered protein synthesis. Importantly, we showed that effects of rG4 in the 5’UTR are context specific and can either increase or decrease protein expression^5^. On one hand, rG4s can promote translation initiation^58^, leading to increased protein levels. On the other hand, rG4s can hinder ribosome scanning^59^, thus decreasing protein expression. These findings reinforce the biological significance of rG4s in translation regulation, supporting the functional importance of rG4s^60^. The significant variant effects observed in *EPN3* and *MSH6* further support G4mer as a strong prediction tool that can inform candidate selection and highlight functionality of genetic variants in the 5’UTR.

Similarly, structural validation using circular dichroism (CD) spectroscopy, a primary tool for the characterization of G4 structures^45^, confirmed G4mer’s predicted structural changes in rG4 formation. The CD spectra demonstrated that the rG4-breaking variant in *EPN3* disrupted the rG4 structure, while the rG4-forming variant in *MSH6* induced the formation of a rG4 structure. However, while the CD spectroscopy results suggest the formation of rG4 structures, we cannot definitively rule out the possibility that these rG4s are forming intermolecularly rather than intramolecularly. This distinction is crucial, as intermolecular rG4s could influence the observed effects differently, potentially impacting the interpretation of their biological significance^61,62^. Nevertheless, these results provide compelling evidence that G4mer can accurately predict the functional and structural impacts of rG4-related variants, further highlighting the critical role that rG4s may play in expression regulation and disease. With these validations, we anticipate that G4mer will enable users who are studying rG4 structures to accurately select sequences and rG4-altering variants for further downstream analyses.

We wish to highlight the following key limitations of the current work. First, unlike our dual luciferase assay experiments, in which the rG4s were surrounded by endogenous flanking sequence, the structure probing experiments lacked this molecular context. Furthermore, we are unable to directly disentangle the potential interplay between sequence, structure, and RBP binding to these elements. As for the training of the G4mer model, we found that different definitions of the training data can significantly impact model performance. While the positive rG4 sequences in this study were derived from rG4-seq experimental protocols, defining the negative non-rG4 sequences presents multiple approaches. In this paper, we enforced a pattern resembling an rG4 in all the negative sequences but were experimentally not detected as rG4s (Methods). Other models have employed different strategies for defining negative sequences, such as shuffling sequences to preserve the dinucleotide distribution of the positive rG4 sequences or including random windows of non-rG4s^63^. Moreover, to define the positively labeled samples in this study, we treated rG4 occurrences as a classification rather than a regression task, therefore potentially limiting the precision in assessing the effects of genetic variants. Thus, future studies could integrate more diverse datasets and consider the design of training data, which could substantially affect the model’s performance and improve generalizability.

We expect that our tool and framework will be valuable for a wide variety of human genetics applications. For instance, G4mer can be applied to analyze ClinVar variants of unknown significance (VUS) or to identify deleterious variants in rare Mendelian diseases. Integrating G4mer scores with additional genomic and clinical data may help to resolve the ambiguity surrounding VUS, a critical need in clinical settings due to the high prevalence of such variants^64,65^. Furthermore, when combined with structural and functional validations, G4mer has the potential to reveal novel disease associations.

The development and application of G4mer underscores the transformative potential of leveraging language models in RNA biology. Given the results presented in this work, combining mRNA language model with functional and structural experimental methods enables us to gain deeper insights into the functional consequences of genetic variants. In line with the performance of language models in RNA research^66–69^, mRNA language models stand as a promising approach to study post-transcriptional mechanisms to identify functionally significant variants that may contribute to disease. As we continue to refine these models and integrate them with experimental validations, we anticipate that such tools will become invaluable in studying variant effects in RNA structures and their implications for human health.

## Methods

### mRNA language model

Recent work surrounding the pre-training and fine-tuning paradigms such as BERT^29^ for natural language processing (NLP), and DNABERT^70^ for the human genome, has shown to perform well for downstream tasks. Following this, to have a model with a general understanding of the sequence structures in the human transcriptome, we developed a pre-trained mRNAbert model. mRNAbert is trained on the sequences in the entire human transcriptome from GENCODE GRCh38, which consists of mature RNA sequences.

To preprocess the sequences, we first removed any duplicates to ensure data integrity. The remaining sequences were then tokenized into overlapping 6-mer tokens, resulting in a maximum input length of 510 tokens per sequence to fit in the 512 token maximum after appending [CLS] and [SEP] tokens. Sequences that resulted in more than 510 6-mer tokens were split into separate inputs without overlaps, since motifs are presumably already captured by the overlapping 6-mer tokens. Hence, this tokenization strategy was chosen to capture local sequence dependencies effectively and potentially learn regulatory motifs in mRNA sequences.

We made slight modifications to the default BERT architecture to better suit the characteristics and data size of mRNA sequences. Specifically, mRNAbert was configured with 6 Transformer layers and 6 attention heads per layer. The model was trained using mixed precision floating point arithmetic to optimize performance and memory usage.

Training was conducted on 4 NVIDIA Tesla P100 GPUs over a period of 3 weeks. The training process involved the following hyperparameters: a learning rate of 4e-4, an effective batch size of 250, and a total of 200,000 training steps. During training, we employed techniques such as gradient accumulation and learning rate scheduling to enhance model convergence and stability.

### rG4 formation and subtype classification datasets

We developed G4mer for rG4 formation prediction by fine-tuning mRNAbert on binary rG4 formation dataset. The preparation of the training dataset includes extracting 5,528 experimentally detected rG4 sequences based on rG4-seq^1^ experimental protocol that have been post-processed and published as rG4-seeker dataset^30^. Duplicated sequences from the same genes that were found by different RT stops were dropped. This left us with 5,454 unique rG4 sequences. Previous work on DNA G4 showed that flanking regions play an important role on the formation and stability of the structure^20^. As such, the rG4 sequences are mapped to the GRCh38.v29 transcriptome sequences to obtain their flanking regions to the left and right of the putative rG4, forming 5,438 unique sequences with a maximum length of 70 nt. Some extended sequences were duplicated when an rG4 is a subset of another rG4, resulting in the same sequence after extension. rG4 sequences that couldn’t be mapped to a transcript were stored as their original sequences without additional flanks.

A similar number of 5,672 negative sequences were obtained by querying all highly expressed genes in the HeLa cell line with no experimentally found rG4s, and looking for a relaxed putative G4 regular expression G(2+)-N(1-30)3-G(2+). The sequences are set to be the same length as the positive sequences, with the matched pattern centered in the sequences. However, since there is a slight over-representation of long patterns found, we set a limit for the number of long patterns included in the training data. We did this by first setting a limit of 1,200 sequences sampled for each bin of 10 nt in the range of 10 nt to 60 nt. The combination of both positive and negative sequences resulted in a total of 11,110 sequences used in the binary dataset.

To develop G4mer for multiclass classification, mRNAbert was fine-tuned on a mutlicass dataset. The 5,454 unique experimentally detected rG4 sequences from rG4-seeker were classified into 8 classes: canonical/G3L1-7, bulges, longloop, two-quartet, G≥ 40%, potential G-quadruplex & G≥ 40%, potential G-triplex & G≥ 40%, and unknown. We obtained their transcriptomic flanking regions when possible to similarly obtain the desired length as for the binary dataset. This resulted in a fine-tuned G4mer model for multiclass classification.

In this work, we sought to explore the potential of mRNA language model and hence included only sequence inputs in the datasets.

### G4mer training and evaluation

For evaluation, we implemented a 10-fold CV strategy to compare the performance of transformer-based G4mer and CNN-based G4Detector. Both models were trained and tested on the same binary and multiclass datasets. The datasets were divided into ten equal parts, with each part used once as the test set while the remaining nine parts were used for training. This ensured that each sequence was tested exactly once.

For the binary classification CV, G4mer was configured with a learning rate of 2e-4, a batch size of 32, and fine-tuned for 2 epochs. G4Detector was run with its default hyperparameters: 256 filters, a kernel size of 12, a batch size of 128, a learning rate of 1e-3, and a hidden size of 32, and was trained for 1 epoch. For the multiclass CV, the hyperparameters for both models remained the same as in the binary classification. However, we implemented early stopping with a patience of 5 for both models, training for a maximum of 50 epochs. The models with the best validation loss were selected for validation. The primary metrics used to assess performance were accuracy and the area under the receiver operating characteristic curve (AUC) for both binary and multiclass classification tasks. The results from all ten folds were averaged to obtain the final performance metrics, providing a robust evaluation of the models’ predictive capabilities.

Using the optimal hyperparameters derived from the CV, G4mer was developed for both binary and multiclass classification. For binary classification, G4mer was trained with a learning rate of 2e-4, a batch size of 32, and trained for 2 epochs using the binary dataset. For multiclass classification, G4mer was trained with the same set of hyperparameters for 5 epochs using the multiclass dataset.

### Model evaluation on G4RNA database

To compare how G4mer performs on sequences from other experimental protocols of various lengths, we extracted sequences from G4RNA. G4RNA sequences were obtained from the G4RNA database^31^. In the data preprocessing phase, duplicate entries were removed from the dataset if they had the same sequence, experimental protocol, and rG4 formation results, eliminating any redundancy that could potentially skew the results of the analyses. Sequences that have disagreeing labels across different experimental protocols, have NaN labels, or appear in the training dataset of G4mer were excluded. The preprocessing step resulted in 795 sequences of lengths ranging from 14 nt to 1368 nt, each validated by one of 24 experimental protocols for rG4 formation.

The final set of G4RNA sequences was used to validate G4mer’s performance. Additionally, we ran 3 other models–cGcC, G4Hunter, and rG4detector–on the same set of sequences. rG4 predictions of G4RNA sequences of cGcC^15^ and G4Hunter^16^ were obtained from G4RNA Screener webtool^63^ that runs both methods in parallel. Default hyperparameters were used, such as the window size of 60 for cGcC and score thresholds of 4.5 for cGcC and 0.9 for G4Hunter. The maximum prediction score per sequence was assigned as the final score. Predictions of rG4detector (https://github.com/OrensteinLab/rG4detector) were obtained by running the tool locally.

### Model interpretation with EIG for flanking sequence importance

To investigate the contribution of rG4 flanking regions to rG4 predictions made by the G4mer model, we utilized EIG for model interpretation. EIG quantifies the importance of individual input features by integrating the gradients of the model’s output with respect to its input, following a path from a baseline class to the input sample (class of interest)^32^. In this context, the baseline serves as a counterfactual reference, reflecting a condition where no rG4 predictive signal is present, while the input sample represents an rG4-containing sequence. By computing and aggregating gradients along a linear or non-linear path between the baseline and the input, EIG provides a detailed attribution score for each feature. For this analysis, our baseline reference is the class of non-rG4 sequences obtained from G4mer’s training data, which serves as the starting point for gradient integration. We then set our input sample as the class of transcriptome-wide predicted canonical rG4 sequences (G4mer score > 0.7) with the rG4 motif centered as much as possible. Since the inputs to G4mer are 6-mers, we obtained an attribution score for every 6-mer input through EIG.

To analyze the k-mer attributions of the rG4 flanking regions, we ran EIG by interpolating between the baseline class of non-rG4 sequences and the input sample class of rG4 sequences, using 200 integration steps. Instead of using a single path, we employed three integration paths, each starting from one of three non-rG4 sequences whose representations are closest to the median representation of all sequences in the baseline class. The final integration path was obtained by averaging the three individual paths. EIG then generates an attribution score for each 6-mer, and we zeroed out the attribution scores for 6-mers that overlapped with the rG4 motifs to focus on flanking regions. We then identified the 6-mers in the rG4 flanking regions that showed significant contributions to the model’s predictions. To assess the significance of these attributions, we conducted a two-sided t-test comparing the k-mer attribution scores of each token and a random set of tokens, adjusting for multiple comparisons using Benjamini-Hochberg procedure with a significance threshold of *FDR* < 0.05 to determine statistical significance.

After obtaining significant 6-mer tokens, we then assigned attribution scores for each nucleotide, as well as 2-mers and 3-mers, by using the maximum attribution score of the significant 6-mer tokens they reside in. To ensure we captured the relevant flanking regions surrounding the rG4 motifs, we extended our analysis to include 20 nucleotides upstream and downstream of the predicted rG4s.

### gnomAD variant analyses

For the analysis of gnomAD variants in UTR regions, we utilized the gnomAD v3.1.2 dataset (https://gnomad.broadinstitute.org/downloads#) for all autosomal chromosomes. Variants were filtered to retain only those that passed all quality control (QC) filters, as indicated by having FILTER value of None. This approach ensures that only high-confidence variants are included in the analysis, reducing the potential impact of sequencing artifacts or low-quality variant calls. We further excluded variants located in low complexity regions, decoy regions, and segmental duplications by checking for the presence of ‘lcr’, ‘decoy’, or ‘segdup’ in the INFO field. These regions are known to introduce biases and inaccuracies in variant calling and are therefore excluded to enhance the reliability of our results. Next, to ensure variants came from reads with high sequencing depth in the gnomAD WGS dataset, only those with a total observed allele count of at least 80% of the maximum number of sequenced alleles (152,312) were retained.

Since transcripts can exhibit varying levels of constraint and selection pressure, we assigned genomic constraints from the gnomAD database (https://gnomad.broadinstitute.org/downloads) to the filtered variants. This allowed us to compare the allele frequency of variants with similar levels of genomic constraints. To achieve this, one group of variants were divided into quartiles based on their genomic constraint scores. For each quartile, we randomly sampled variants from the other group. The stratified sampling ensured that both groups contained variants had closely matched quartile distributions.

To analyze rG4-breaking variants using the CADD metric, we retrieved raw CADD scores for each variant from the gnomAD Hail Table, which assigns a raw CADD score for every variant based on its predicted deleteriousness. Specifically, we intersected our list of rG4-breaking variants with the gnomAD data, matching each variant by its locus, reference allele, and alternate allele. The matched raw CADD scores reflect the relative likelihood that a given variant is deleterious based on a wide range of annotations, including sequence conservation and regulatory impact where a positive CADD score represents increased potential deleteriousness. The scores were then used in the analyses to assess the potential functional impact of variants that disrupt rG4 structures. For the rG4-breaking variant in 5’UTR, we obtained the raw CADD scores for all variants. For 3’UTR, we focused on rG4-breaking variants in disease genes, particularly those with LOEUF upper bound values < 0.9 from the gnomAD v4.1 transcript constraint file.

### Penn Medicine BioBank

Penn Medicine BioBank (PMBB) hosts a comprehensive median of seven years of longitudinal electronic health records (EHR), genetic sequencing, and biological samples of study participants. Participants for the PMBB study were enlisted from clinical practice sites of the University of Pennsylvania Health System. Each participant gave informed consent for the retention of their health data and utilization for research purposes, including permission to be potentially recontacted for future research endeavors. The study was also sanctioned by the University of Pennsylvania’s Institutional Review Board, ensuring adherence to the ethical guidelines outlined in the Declaration of Helsinki. The PMBB study’s dataset encompasses a subgroup of over 44,000 individuals who underwent Whole Exome Sequencing (WES). DNA was isolated from the stored buffy coats of these individuals, and exome sequencing was carried out by the Regeneron Genetics Center in Tarrytown, NY, aligning the sequences with the GRCh38 reference genome as previously outlined^28^. In preparation for further phenotype analysis, samples were excluded based on criteria such as low exome sequencing coverage (below 85% of targets reaching 20x coverage), high rates of heterozygosity/contamination (D-stat > 0.4), genetically identified sample duplicates, and discrepancies between reported and genetically verified sex.

We further filtered samples and variants in PMBB to be included in our analyses. We restricted the analyses to only keep samples who are 2nd degree unrelated in each ancestry group, resulting in two groups of samples of European (N=29,362) and African (N=10,217) genetic ancestries. Furthermore, exclusion criteria applied to variants include singletons and high missing call rates (exceeding 0.1).

To obtain PMBB variants that are rG4-altering, we mapped variants across all transcripts and obtained the wild-type regions around the variants. rG4-breaking variants were defined by a wild-type G4mer score above the rG4 threshold of 0.5, with the mutated sequence score at least 0.2 lower than the wild-type to indicate reduction in rG4 structure stability in the presence of the variant. Conversely, rG4-forming variants were characterized by a wild-type score below 0.5, indicating the absence or weak formation of rG4, and a mutated score at least 0.2 higher, suggesting increased rG4 stability or formation in the presence of the variant.

### Phenome-wide association studies

Variants found in PMBB dataset were assessed for their effects on putative rG4s found in transcripts. We tested association of these variants with phenotypes extracted from PMBB^28^. ICD-10 codes of samples from PMBB were mapped to distinct disease entities (i.e. phecodes) via Phecode Map 1.2 using the PheWAS package in R^71^. To establish a patient as a ‘case’ for a given phecode, we required a minimum of 2 counts of the code, as repeated diagnoses of a code on different days improves phenotype precision^72^. Our association analyses considered only phecodes with at least 20 cases based on prior simulation studies for power analysis of common and rare variant gene burden PheWAS on binary traits, highlighting the influence of the number of cases on the power to detect genetic associations^73,74^. This led to the inclusion of 957 phecodes. Each phecode was tested for association with each rG4-altering variant using a logistic regression model adjusted for sex, age at enrollment, age-squared, and the first 10 principal components of genetic ancestry. For each variant, ancestry-specific PheWAS was first performed for two major groups: African (AFR) and European (EUR) ancestry. The cross-ancestry summary-results were then meta-analyzed using an inverse-variance weighted random-effects model^28^. The odds ratio (OR) from the meta-analysis of combined ancestry is used to determine the direction of each arrowhead in the association plots, where OR *>* 1 suggest an increased risk with an up arrowhead.

To address multiple testing, an association between variant and phecode with FDR p < 0.1 was considered significant. Thus, the adjusted threshold for significance was p « 5.53e-2. In addition, we calculated a more stringent Bonferroni corrected threshold for each gene of p « 9.78e-5. We then filtered the rG4-altering variants for functional validation using dual luciferase assays by considering those with associations that pass the Bonferroni adjusted significance threshold.

### Plasmid preparation, cell culture, and transfections

In order to perform a bicistronic dual luciferase assay to quantify changes in expression due to variants in predicted rG4s, we created a modified version of pMiRcheck2^75^, puORF-Check3 (Supplementary Fig. S5). We obtained the WT 5’UTR sequences of *EPN3* and *MSH6* (387 and 188 nt respectively), and the first 12 nt of CDS, to clone upstream of the stuffer sequence in the puORF-Check3 plasmid backbone using the NdeI restriction site (NEB HiFi). Mutated sequences of each 5’UTR were cloned in the same manner as well. All constructs were verified by whole plasmid sequencing. HuH-7, NIH3T3, and CHO cells were used for conditional expression of reporter genes. For transient transfections, cells were seeded 1 day before transfection in 24-well plates at a density of 80,000 cells per well. 60 ng of the dual luciferase plasmid was transfected into each well using Lipofectamine 3000 following the manufacturer’s protocol, with 3-4 technical repeats for each construct. 3-5 biological replicates were obtained by transfecting cells from separate passages on separate days using newly prepared reagents. HuH-7 and NIH3T3 cells were cultured in Dulbecco’s modified Eagle’s medium (DMEM), and CHO cells were cultured in F12 medium, both supplemented with 10% fetal bovine serum.

### Dual luciferase reporter assays

Luminescence was measured using the Promega Dual-Luciferase Reporter Assay System (E1910) following the manufacturer’s protocol. Cells were lysed by adding 100 *µ*L of lysis buffer, and 20 *µ*L of each lysate were transferred to a white opaque 96-well plate. With Renilla luciferase serving as the reporter and Firefly luciferase as the internal control, the intensities of Firefly and Renilla luciferase luminescence were measured with a microplate reader (BioTek Synergy Neo2 multimode reader) after automatic injection (BioTek Dual Reagent Injector by Agilent) of the Luciferase Assay Reagent II and Stop & Glo reagents. For each test construct, the Firefly and Renilla luciferase activities were inferred from the measured counts per minute to obtain the relative Firefly-to-Renilla expression ratio. Measurements for mutated constructs were then normalized to the wild-type construct to provide a fold-change measure for the translational efficiency conferred by each variant, as described in previous studies of translational regulation^76^. Statistical significance was determined using independent t-test, comparing the relative Firefly-to-Renilla expression ratios across transfections of each wild-type and mutated construct pair for all cell lines.

### Circular dichroism (CD) spectra

RNAs were purchased from Integrated DNA Technologies (IDT) and prepared as follows: RNA stock solutions were diluted to a final concentration of 40 µg/mL in buffer containing 150 mM KCl or 150 mM LiCl. Prior to analysis, all RNA samples were heated to 100°C for 3 minutes and then allowed to cool to room temperature.

Circular dichroism (CD) spectra were collected using a Chirascan V100 spectrometer over a wavelength range of 220-300 nm, with a step size of 0.5 nm, a bandwidth of 1 nm, and an integration time of 1.25 seconds per data point. Spectra were recorded at 25°C for initial measurements. Following the initial spectra collection, the sample chamber temperature was increased to 95°C using a Peltier controller, and the samples were allowed to stabilize for 3 minutes before additional spectra were acquired. Baseline spectra of the buffer (150 mM KCl or 150 mM LiCl) were subtracted from the RNA spectra using the Chirascan software to ensure accurate measurements. All CD spectra were obtained in three independent replicates under each condition.

## Acknowledgements

We sincerely thank Matthew Gazzara, Seong Woo Han, Kevin Yang, David Wang, and Di Wu for their contributions and suggestions. Our appreciation extends to the members of the Barash lab for discussions and thoughtful feedback. Special thanks to Michelle Scott and Jean-Michel Garant (Sherbrooke University) for sharing sequences in G4RNA database. Finally, we gratefully acknowledge the staff of the Regeneron Genetics Center for whole-exome sequencing of PMBB participants. This work was supported by NIGMS grants T32GM148376 (to A.J.), R35GM142864 (to D.D.), and R25GM055366 (to B.G.); NHGRI grant T32HG009495 and The Zuckerman-CHE STEM Leadership Program (to D.G.); CureBRCA and the Basser Center for BRCA pilot grants (to Y.B. and K.N.); and NIH grants R01 LM013437 and GM-147739 (to Y.B.).

## Author contributions statement

F.Z. and Y.B. conceived and designed the project. F.Z. developed mRNAbert and G4mer, performed model performance analyses, variant analyses, and disease association studies with ancestry-based PMBB variants under the guidance of Y.B. PMBB samples and variants for disease association analyses filtering were done by F.Z and D.G under the guidance of Y.B. Selection of disease-associated variants and sequences for validation were done by F.Z. and D.G under the guidance of Y.B. and K.N. N.I. and F.Z. performed EIG analyses under the guidance of Y.B. N.H designed puORF-Check3. D.G. planned and performed the cloning and dual luciferase experiments and analyses under the guidance of Y.B and N.H. Selection of sequences for structural validations were done by F.Z. under the guidance of Y.B. and D.D. A.J. and B.G. performed circular dichroism experiments and analyses under the guidance of D.D. D.G. provided the methods on dual luciferase experiments. A.J. provided methods on circular dichroism experiments. F.Z. and S.J. designed the web tool and S.J. built the web tool under the guidance of Y.B. F.Z. wrote the paper and all authors contributed to editing the paper.

