## Supplementary Information for "G4mer: An RNA language model for transcriptome-wide identification of G-quadruplexes and disease variants from population-scale genetic data"


#### CONTENTS


**1 Supplementary Figures** **2**



### 1. SUPPLEMENTARY FIGURES

a

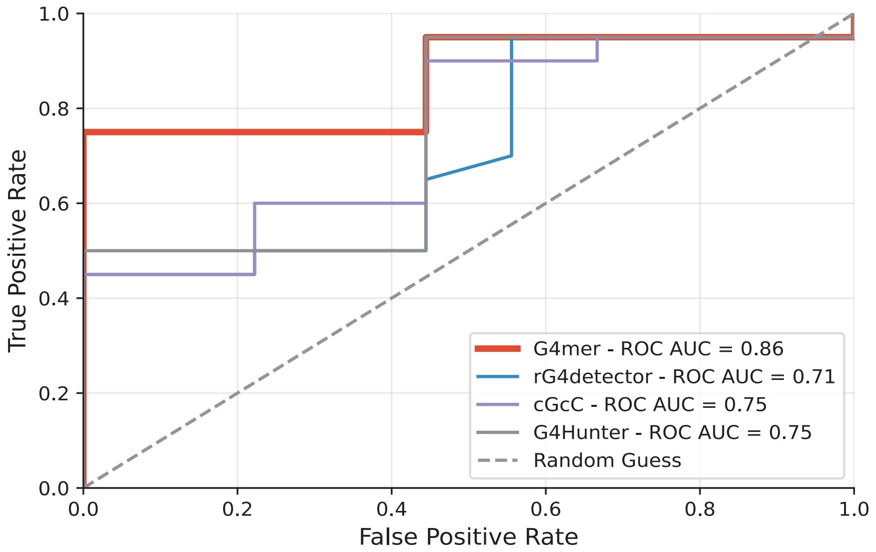


b

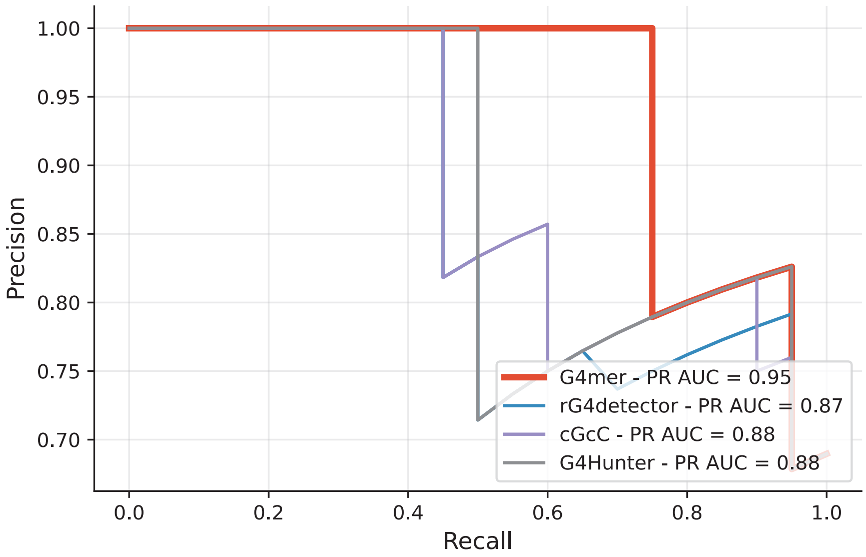


**Fig. S1. Model performance comparison on sequences from G4RNA database.** **(a)** ROC curves and **(b)** Precision-Recall (PR) curves comparing the performance of G4mer, rG4detector, cGcC, and G4Hunter in predicting rG4 formation for sequences from G4RNA database. Sequences were validated by various experimental protocols.

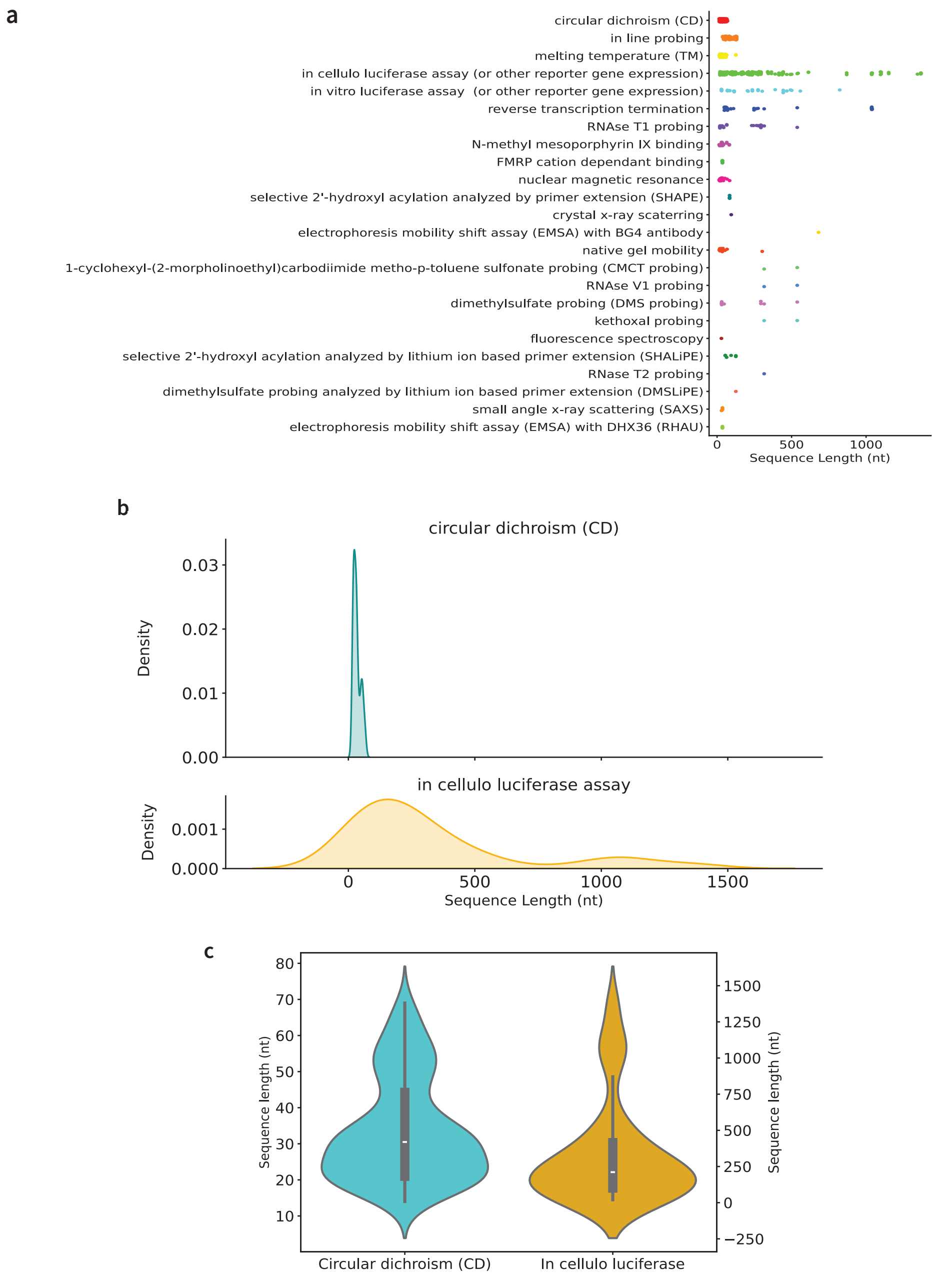


**Fig. S2. G4RNA database sequence length distribution per experimental protocol.** **(a)** Lengths of sequences tested by each of the 24 experimental protocols in G4RNA **(b)** Density plots showing the distribution of sequence lengths (in nucleotides) for two experimental protocols: circular dichroism (CD) and in cellulo luciferase assay. The top panel illustrates the density of sequence lengths for circular dichroism, where the majority of sequences are clustered in a narrow length range below 100 nt. The bottom panel shows the distribution for in cellulo luciferase assay, where the sequence lengths are more widely dispersed, covering a broader range of values with the longest sequence being 1,368 nt. **(c)** Violin plots comparing the distribution of sequence lengths between circular dichroism (CD) and in cellulo luciferase assay. The left y-axis corresponds to the sequence lengths for CD, where the median is 30.5 nt as indicated by the horizontal white line within the violin plot. The right y-axis provides a scale for the in cellulo luciferase assay's broader sequence length range where the median is 210 nt.

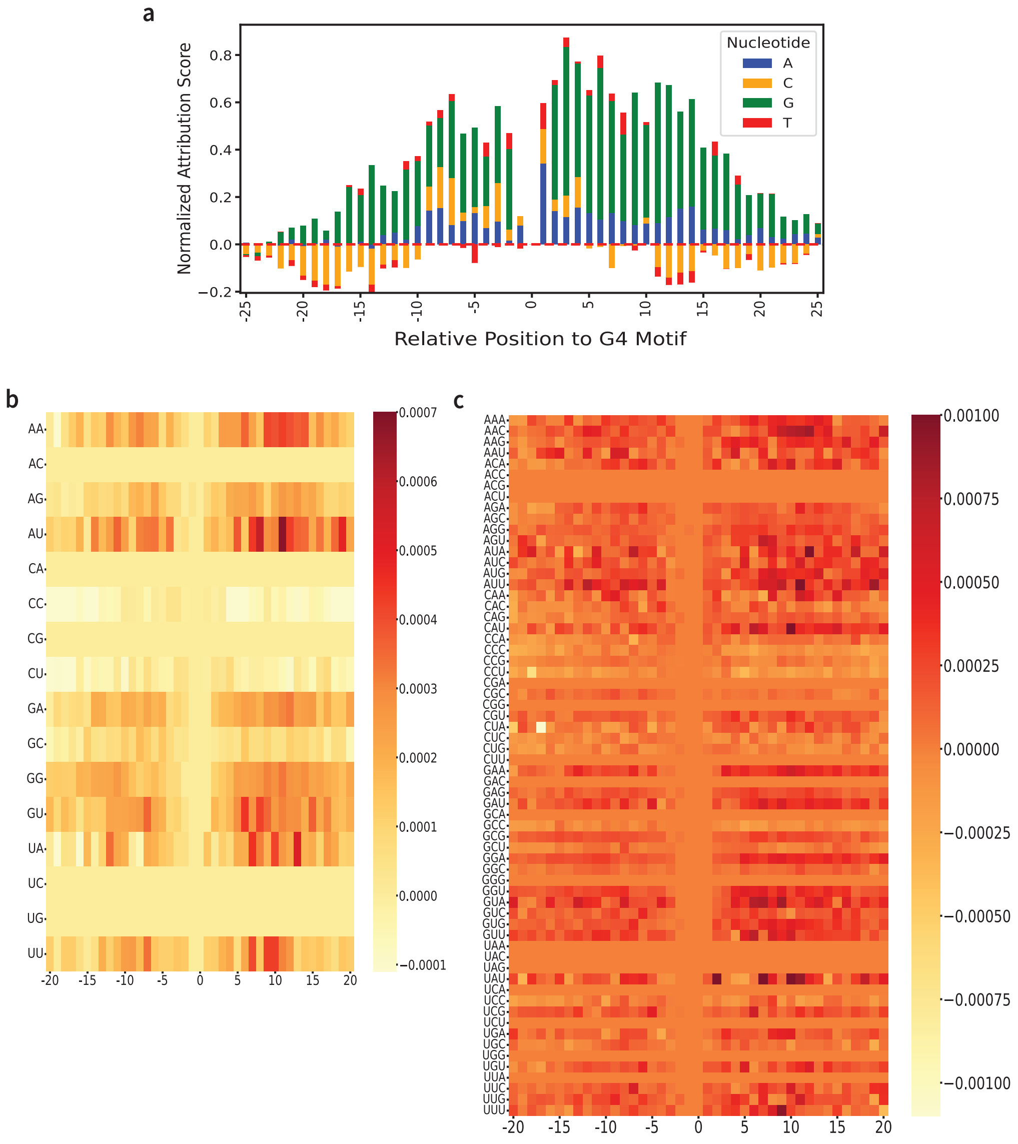


**Fig. S3. Attribution scores of rG4 flanks from EIG analysis. (a)** Stacked bar plot showing normalized attribution scores for individual nucleotides (A, C, G, T) at each position relative to the rG4 motif across G4mer predicted rG4 sequences transcriptome-wide. The x-axis represents positions upstream and downstream of the G4 motif, with 0 corresponding to the rG4 motif. Positive scores indicate a stronger contribution to rG4 formation, while negative scores suggest reduced influence. **(b, c)** Heatmaps of normalized attribution scores for **(b)** 2-mers and **(c)** 3-mers across positions relative to the G4 motif. Each row represents a unique k-mer, and each column corresponds to a position relative to the G4 motif, with color intensity indicating the magnitude of the attribution score and higher scores indicating stronger contribution to rG4 structure prediction by G4mer.

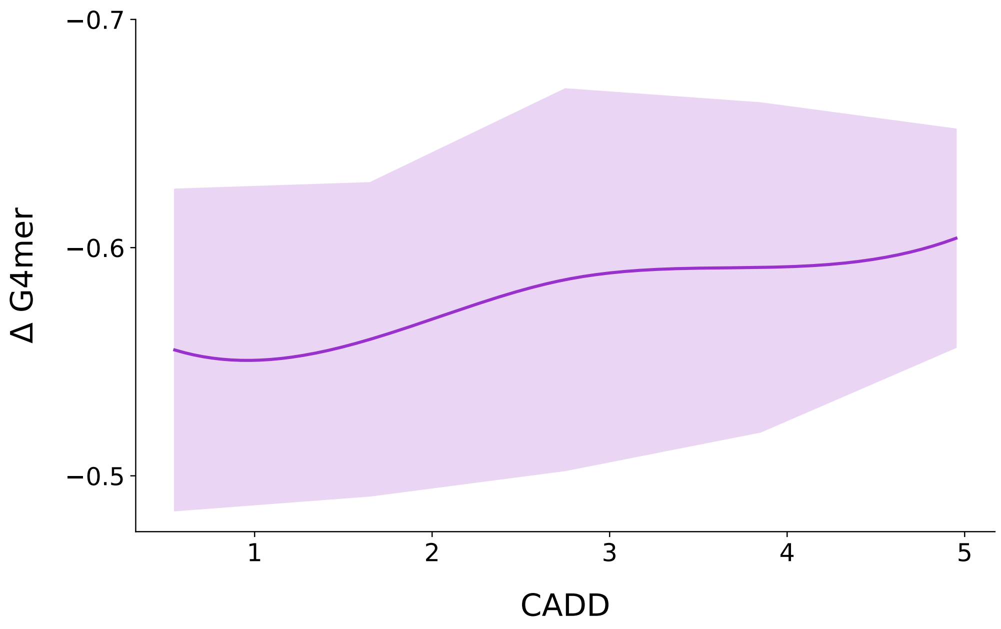


**Fig. S4. Relationship between CADD scores and ΔG4mer values in the 3'UTR.** The plot shows the relationship between the Combined Annotation Dependent Depletion (CADD) score and the change in G4mer (ΔG4mer) values in the presence of rG4-breaking variants in disease genes. The purple line represents the mean ΔG4mer value across different CADD scores, while the shaded area indicates the standard deviation. As the CADD score increases from 1 to 5, the ΔG4mer values display a subtle upward trend, suggesting a slight increase in deleteriousness of the variants that have greater breaking effects on rG4 structures in the 3'UTR regions.

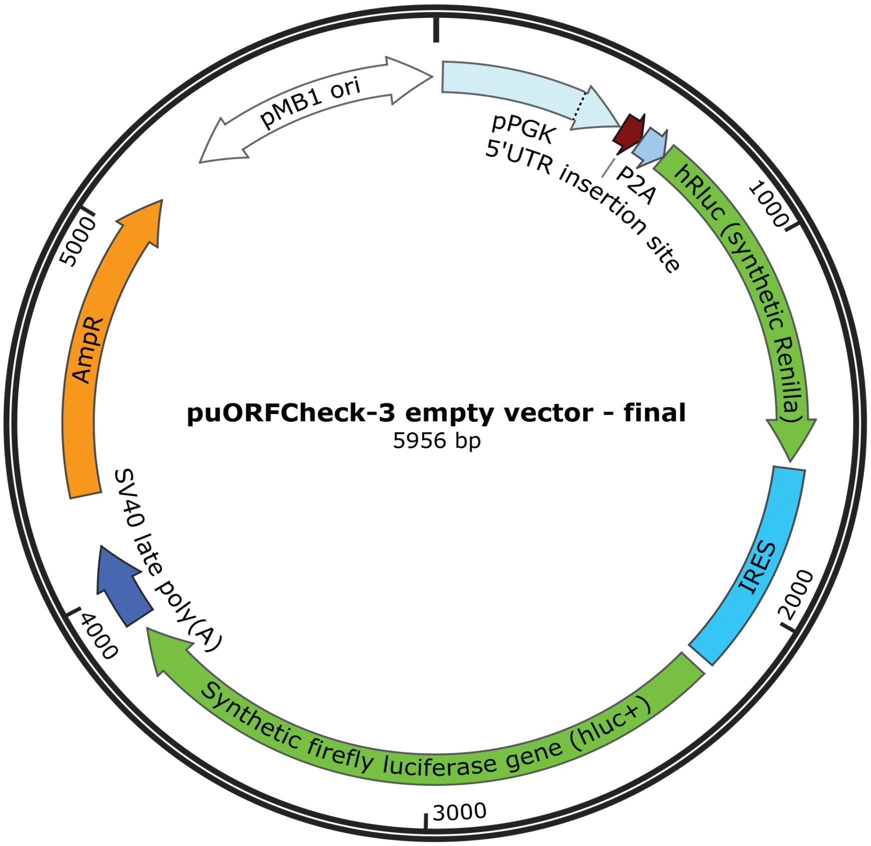


**Fig. S5. puORF-Check3 plasmid map for dual luciferase reporter assays.** The plasmid puORF-Check3 is a 5.9 kb vector designed for mammalian expression of a bicistronic dual-luciferase system. It features a pMB1 origin of replication (pMB1 ori) to ensure efficient plasmid replication in E. coli. A phosphoglycerate kinase (pPGK) promoter, located upstream of the NdeI cloning site (labeled as the 5'UTR insertion site), drives high-level expression in mammalian cells. A P2A self cleaving peptide is fused to the hrluc+ luciferase sequence to ensure 5'UTR frame is conserved. The plasmid encodes two luciferase genes, hRluc (Renilla luciferase) and hluc+ (Firefly luciferase), transcribed together as a single mRNA transcript. These genes are separated by a viral internal ribosomal entry site (IRES), enabling simultaneous measurement of luminescence from both luciferase proteins in cell lysates. Additionally, an ampicillin resistance gene (AmpR) is included to facilitate antibiotic selection in E. coli and allow for the propagation of successfully transformed cells.
